# High Phylogenetic Utility of an Ultraconserved Element Probe Set Designed for Arachnida

**DOI:** 10.1101/065201

**Authors:** James Starrett, Shahan Derkarabetian, Marshal Hedin, Robert W. Bryson, John E. McCormack, Brant C. Faircloth

**Author notes:** equal contribution (James Starrett, Shahan Derkarabetian). Corresponding author: Shahan Derkarabetian, Department of Biology, San Diego State University, 5500 Campanile Dr., San Diego, CA 92182-4614.

## Abstract

Arachnida is an ancient, diverse, and ecologically important animal group that contains a number of species of interest for medical, agricultural, and engineering applications. Despite this applied importance, many aspects of the arachnid tree of life remain unresolved, hindering comparative approaches to arachnid biology. Biologists have made considerable efforts to resolve the arachnid phylogeny; yet, limited and challenging morphological characters, as well as a dearth of genetic resources, have confounded these attempts. Here, we present a genomic toolkit for arachnids featuring hundreds of conserved DNA regions (ultraconserved elements or UCEs) that allow targeted sequencing of any species in the arachnid tree of life. We used recently developed capture probes designed from conserved genomic regions of available arachnid genomes to enrich a sample of loci from 32 diverse arachnids. Sequence capture returned an average of 487 UCE loci for all species, with a range from 170 to 722. Phylogenetic analysis of these UCEs produced a highly resolved arachnid tree with relationships largely consistent with recent transcriptome-based phylogenies. We also tested the phylogenetic informativeness of UCE probes within the spider, scorpion, and harvestman orders, demonstrating the utility of these markers at shallower taxonomic scales, even down to the level of species differences. This probe set will open the door to phylogenomic and population genomic studies across the arachnid tree of life, enabling systematics, species delimitation, species discovery, and conservation of these diverse arthropods.

## Introduction

Arachnida is an extremely ancient and diverse arthropod lineage, including conspicuous taxa such as spiders, scorpions, mites, and ticks. The oldest known arachnid fossils consist mostly of scorpions and extinct trigonotarbids from the Silurian period, 443-416 million years ago (MYA; Laurie 1899; Jeram *et al*. 1990; Dunlop, 1996; Dunlop et al. 2008). More than 110,000 species of arachnids have been described, with spiders and mites ranking among the most diverse of all animal orders (Harvey 2002; Zhang 2011). Yet, more than half of arachnid species diversity remains to be discovered (Chapman 2009). Arachnida also contains a number of medically important venomous or disease-vector species and many important agricultural mite pests, while spiders are of particular interest to biologists and engineers for the strong and elastic silk fibers they produce. Despite the attention arachnids have received for their ecological importance and practical utility to humans, phylogenetic relationships among and within many arachnid orders remain uncertain. At the root of this problem is that morphological characters are limited and difficult to interpret (Shultz 2007), and genomic resources for many species in this group are sparse. Adding to the difficulty is the uncertainty in the rooting of the arachnid tree, with fossil, morphological, and molecular data recovering drastic discrepancies in early arachnid relationships and placing traditional non-arachnid chelicerates within Arachnida (Wheeler and Hayashi 1998; Masta *et al.* 2009; Regier *et al.* 2010; Sharma *et al.* 2014). Genome wide phylogenetic markers are essential for resolving deep relationships within Arachnida and for helping to uncover the numerous arachnid species that await discovery.

Ultraconserved elements (UCEs) provide one potential target for a universal set of genomic markers for arachnids and would allow researchers to collect genomic information from diverse taxa across the arachnid tree of life. UCEs are segments of DNA that are highly conserved across divergent taxa (Bejerano *et al.* 2004) and are thought regulate and/or enhance gene expression (Alexander *et al.* 2010). UCEs can also be used as anchors to target and retrieve variable DNA sequence flanking the core UCE regions, and the flanking DNA shows a trend of increasing genetic variability as distance from the core UCE increases (Faircloth *et al.* 2012). UCEs are ideal markers for molecular systematics for several reasons. Whereas transcriptomes require high-quality RNA as input, the UCE protocol only requires DNA, and enrichments can be performed using relatively low starting DNA concentrations. This allows the method to be extended to small-bodied taxa, large collections of specimens preserved for Sanger-based DNA research, and even to “standard” museum specimens with varying levels of DNA degradation (McCormack *et al.* 2015). Homology between UCE loci across divergent taxa is also easy to assess because the core UCE region often displays >95% sequence similarity, and UCE cores are rarely duplicated or paralogs (Derti *et al*. 2006). Although core UCE regions show reduced sequence variation, UCE-flanking DNA shows levels of phylogenetic informativeness equal to or greater than that of traditionally used protein-coding markers (Gilbert *et al.* 2015).

UCEs have been successfully used in phylogenetic studies at multiple taxonomic levels, including shallow, young divergences (~5 MYA, within species; Smith *et al.* 2014; Harvey *et al.* 2016; Manthey *et al.* 2016) and deep, ancient divergences (e.g., all amniotes; Faircloth et al. 2012). Originally developed for use in tetrapods, the vast majority of UCE phylogenetic studies have been conducted on vertebrate taxa including all amniotes (Faircloth et al. 2012), mammals (McCormack *et al.* 2012), birds (McCormack *et al.* 2013; Smith *et al.* 2014), reptiles (Crawford *et al.* 2012; Crawford *et al.* 2015), and fish (Faircloth *et al.* 2013). More recently, UCE probe sets have been developed for use in arthropod taxa, including Hymenoptera, Coleoptera, Diptera, Lepidoptera, and Hemiptera (Faircloth *et al.* 2015; Faircloth 2016).

Here, we perform an *in vitro* test of a recently developed bait set targeting arachnid UCEs (Faircloth 2016). We then demonstrate the phylogenetic utility of this probe set at multiple evolutionary timescales, from those spanning hundreds of millions of years to species-level divergences of <10 million years. We reconstruct well-supported phylogenies between and within arachnid orders, and we demonstrate that these UCE baits may also be useful when reconstructing species level relationships.

## Materials and Methods

### UCE library construction and enrichment

UCE capture was tested using the master arachnid bait set (Faircloth 2016) on 32 arachnid samples representing six orders (Table 1). The orders selected span the root of Arachnida (Regier *et al.* 2010; Sharma *et al.* 2014). Within three orders (Araneae, Opiliones, Scorpiones), taxa were selected from major lineages, including across the internal root for each (Hedin *et al.* 2012; Bond *et al.* 2014; Sharma *et al.* 2015). To assess UCE variability between closely related taxa, we included two turret spiders from the *Antrodiaetus riversi* complex (Hedin *et al.* 2013), two harvestmen representing *Briggsus pacificus* and *B. bilobatus*, and a second bark scorpion *Centruroides sculpturatus* to complement the published *C*. *exilicauda* genome. Voucher specimens are located in the San Diego State University Terrestrial Arthropod Collection (SDSU_TAC). In the case of small arachnids (e.g., mites and ticks), whole specimens were used in extractions, but vouchers from the same locality are deposited in SDSU_TAC.

**Table 1.**
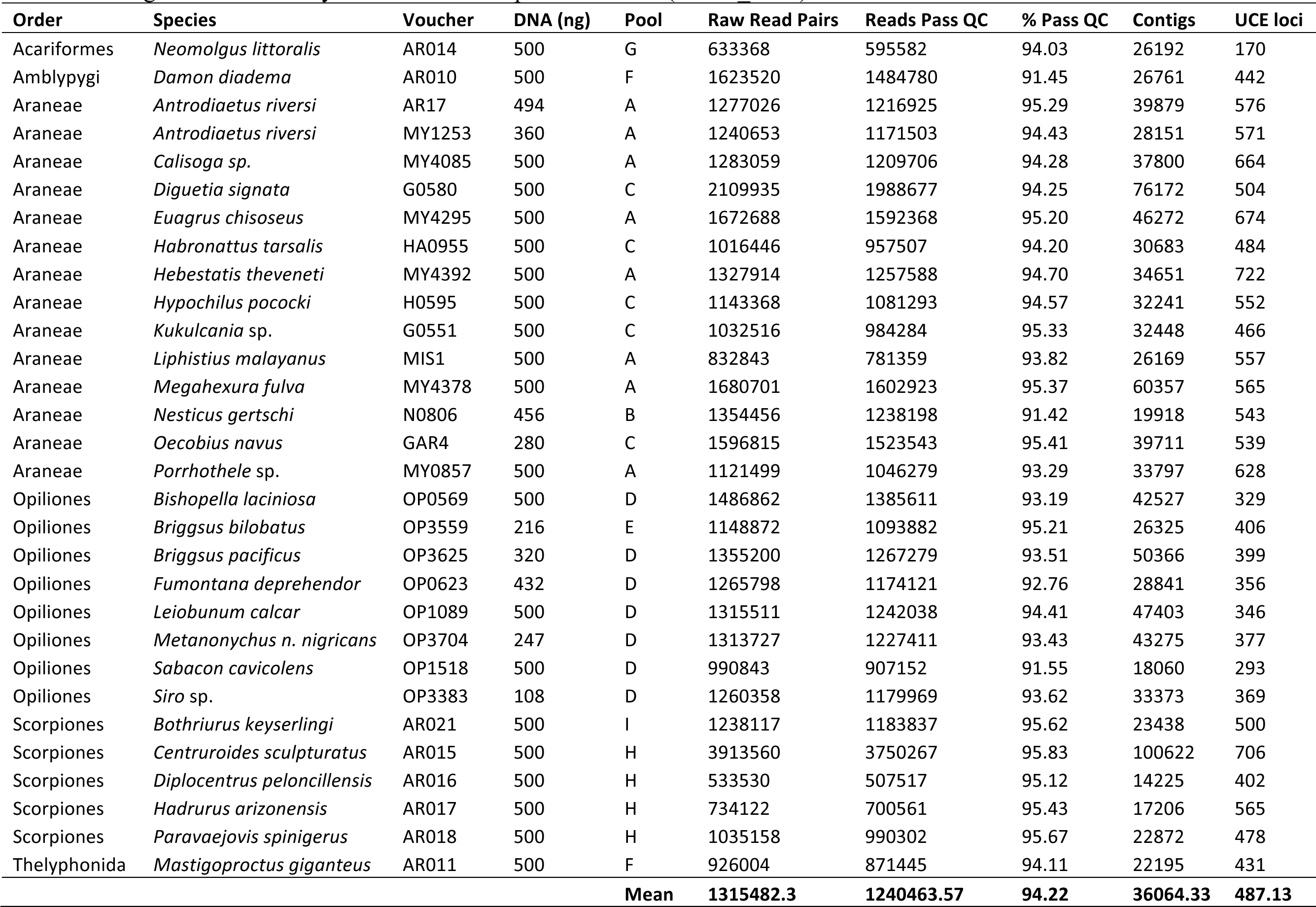
Sample information and sequencing statistics. DNA refers to starting quantity that was processed. Voucher specimens are housed in the San Diego State University Terrestrial Arthropod Collection (SDSU_TAC).

Genomic DNA was extracted from legs or whole specimens using the DNeasy Blood and Tissue Kit (Qiagen, Valencia, CA) following the manufacturer’s protocol. DNA concentrations were determined with a Qubit fluorometer (Life Technologies, Inc.) and run out on a 1% agarose gel to assess quality. The lowest starting DNA quantity for processing was 108 ng (*Siro acaroides*, a small-bodied harvestmen typically <2mm in length), while most samples started with approximately 500 ng (Table 1). Samples with high molecular weight DNA were fragmented with a QSonica Q800R sonicator for 10–12 cycles of 20 sec on/20 sec off, resulting in fragments predominantly in the range of 300–1000 bp. DNA from sample AR014 ( *Neomolgus littoralis*) was partially degraded and not sonicated.

Libraries were prepared with the KAPA Hyper Prep Kit (Kapa Biosystems), with a generic SPRI substitute used for bead clean-up steps (Rohland and Reich 2012; Glenn *et al.* 2016). Universal adapters were ligated onto end-repaired and A-tailed DNA fragments. Each adapter-ligated library was amplified in a 50 µl total reaction, which consisted of 15 µl of adapter ligated DNA, 1X KAPA HiFi HotStart ReadyMix, and 5 µM of each Illumina TruSeq dual-indexed primer (i5 and i7) with modified 8 bp indexes (Glenn *et al.* 2016). Amplification conditions were 98°C for 45 sec, followed by 10-12 cycles of 98°C for 15 sec, 60°C for 30 sec, and 72°C for 60 sec, and then a final extension of 72°C for 60 sec. After bead clean-up of amplified libraries, equimolar amounts of libraries were combined into 1000 ng total pools. Pool combinations ranged from 1–8 individual libraries (Table 1). Due to a low concentration following library preparation and amplification, the pool containing sample AR014 ( *Neomolgus littoralis*) included a lower amount of library (108.9 ng) compared to the other two libraries that we included (333 ng each).

Target enrichment of libraries was performed using the MySelect kit (Microarray) following the Target Enrichment of Illumina Libraries v. 1.5 protocol (http://ultraconserved.org/#protocols). Custom TruSeq adaptor blockers (Glenn *et al.* 2016) and standard MYbaits blockers were annealed to 147 ng/uL library pools, followed by hybridization to the master arachnid bait set (Faircloth 2016). Hybridizations were performed at 65°C for 24 hours. After hybridization, library pools were bound to Dynabeads MyOne Streptavidin C1 magnetic beads (Life Technologies) for enrichment. We performed with-bead PCR recovery of the post-hybridization enrichments in a 50 µl volume reaction consisting of 15 µl enriched DNA, 1X KAPA HiFi HotStart ReadyMix, and 5 µM each of TruSeq Forward and Reverse primers. Amplification conditions were 98°C for 45 sec, followed by 18 cycles of 98°C for 15 sec, 60°C for 30 sec, and 72°C for 60 sec, and then a final extension of 72°C for five minutes. Following PCR recovery, libraries were quantified using a Qubit fluorometer and diluted to 5 ng/µL. We performed qPCR quantification of enriched library pools, and we prepared a 10 uM mix of each pool at equimolar ratios. We sequenced the library pool using a partial run of paired-end bp sequencing on an Illumina NextSeq (Georgia Genomics Facility).

### Read Processing, Contig Assembly, and Matrix Creation

Raw read data were processed using the PHYLUCE pipeline (Faircloth 2015). Adapter removal and quality control trimming were done using the Illumiprocessor wrapper (Faircloth 2013) using default values. Reads were assembled using Trinity version r2013-02-25 (Grabherr *et al.* 2011). Contigs from all samples were matched to probes using minimum coverage and minimum identity values of 65. We additionally extracted UCE loci *in silico* from available arachnid genomes and *Limulus polyphemus*. UCE loci were aligned using mafft (Katoh and Standley, 2013) and trimmed with gblocks (Castresana 2000; Talavera and Castresana 2007) as implemented in the PHYLUCE pipeline. Multiple datasets were created for downstream analyses. First, a “2perArachnid” dataset was created that contained up to two representative samples from each arachnid order sequenced, plus *Limulus*as an outgroup. This included UCE data extracted from both novel samples and previously deposited genomes. Representative samples were chosen to span the root node of the relevant arachnid orders and based on number of UCE loci recovered. Because the purpose of our study was not to reconstruct arachnid phylogeny, we did not include all arachnid orders. Second, we assembled a “UCEsample” dataset that included only samples newly sequenced for this study. Third, three individual datasets were created that included all samples from within the orders Araneae, Opiliones, and Scorpiones. For each of these individual-order datasets, two matrices were created, one including the amblypygid *Damon* as the outgroup, and a second without an outgroup. Data matrices without outgroups were used for determining matrix statistics, with the number of parsimony informative characters computed using PAUP* 4.0 (Sinauer Associates, Inc.). Finally, to assess species-level utility of UCEs, three congeneric datasets were created for *Antrodiaetus* (turret spiders), *Briggsus* (Briggs’ harvestmen), and *Centruroides* (bark scorpions).

### Phylogenetic Analysis

Data matrices including an outgroup taxon were also subjected to phylogenetic analyses using RAxML HPC v8.0 (Stamatakis 2014), implementing the rapid bootstrap algorithm (Stamatakis *et al.* 2008) plus ML tree search option (-f a), 200 bootstrap replicates, and the GTRGAMMA model. ML analyses were conducted on two matrices for each dataset, one with 70% taxon coverage for each locus and one with 50% coverage. Topologies and support scores based on 70% taxon coverage were nearly identical to those from 50% taxon coverage, and we show matrix statistics and trees for only 50% datasets. All analyses were conducted on a late 2015 iMac with a 4GHz Intel i7 processor, with the exception of contig assembly with Trinity, which was run on a 12 core CentOS linux machine with 48 GB of RAM.

## Results

Sequencing results, assembly statistics, and number of UCE loci recovered are presented in Table 1 for samples sequenced *in vitro* in this study and in SuppTable 1 for published genomes used *in silico*. On average, we produced 1,315,482 raw reads per sample, with an average of 1,240,464 reads (94.2%) passing quality control. Assemblies resulted in an average of 35,884 contigs per sample. The number of UCE loci recovered from newly sequenced samples varied between 170 and 722 (average=487), while the average recovered *in silico* from published genomes was 675, including 555 UCEs from the outgroup *Limulus*. The average number of recovered UCE loci differed (unpaired two tailed t-test, t = 6.4308, df = 38, p-value = <0.0001) among orders, being higher in groups from which the probes were designed (Araneae, Parasitiformes, Scorpiones: 605 loci) versus those that were not used in probe design (Acariformes, Amblypygi, Opiliones, Thelyphonida: 376 loci). We tested whether the number of samples included in a hybridization pool influenced the number of UCE loci recovered and found no correlation (R^2^ = 0.03), although the Opiliones-only pools recovered the fewest loci (SuppFigure 1). Matrix statistics are presented for a 50% taxon coverage dataset in Table 2. We recovered identical topologies and nearly identical support scores for matrices with 70% taxon coverage (not shown) despite an approximate 37% decrease in loci numbers. Across the final matrices, an average of 589 UCE loci were included in the 50% matrices, the lowest being the Opiliones + OUT datasets. Matrix lengths varied from 105,337 bp in the “UCEsample” matrix to 452,309 bp in the Scorpiones matrix. The average percentage of parsimony-informative characters was 28.99%.

**Table 2.** Matrix statistics. PI = parsimony informative sites.

| Dataset | N | 50% taxon coverage |  |  |  |  |  |
| --- | --- | --- | --- | --- | --- | --- | --- |
|  |  | Loci | Length | Mean Length | Min-Max length | PI | % PI |
| 2per Arachnid | 11 | 602 | 199,810 | 331.91 | 113-853 | 60,468 | 30.3 |
| UCEsample | 32 | 510 | 105,337 | 206.54 | 80-746 | 49,362 | 46.9 |
| Araneae +OUT | 20 | 686 | 169,134 | 246.55 | 92-833 | 67,644 | 40.0 |
| Araneae | 19 | 724 | 181,915 | 251.26 | 92-833 | 71,786 | 39.5 |
| Opiliones +OUT | 9 | 435 | 158,314 | 363.94 | 97-893 | 33,263 | 21.0 |
| Opiliones | 8 | 381 | 153,187 | 402.07 | 97-914 | 29,807 | 19.5 |
| Scorpiones+OUT | 8 | 627 | 318,178 | 507.46 | 145-1136 | 65,227 | 20.5 |
| Scorpiones | 7 | 749 | 452,309 | 603.88 | 99-1141 | 64,892 | 14.4 |
| **Mean** |  | **589.3** | **217273** | **364.2** |  | **55306.1** | **29.0** |

For shallow time-scale comparisons, *Briggsus* (Briggs’ harvestmen) resulted in the lowest number of UCE loci recovered (Table 3; 292 loci, total length of 172,562, average locus length 591), but contained the highest number of variable sites (25,524 - 14.8%). Conversely, the *Centruroides* (bark scorpions) comparison recovered the highest number of UCE loci (585 loci, total length of 583,454, average locus length 997.4) but lowest number of variable sites (7,660 - 1.3%). The proportion of variable loci for all comparisons were above 0.9, with average number of variable sites per locus ranging from 14–87 (Table 3).

**Table 3.** Congeneric matrix statistics.

| Dataset | Loci | Total Length | Mean Length | Min-Max length | Variable Sites | % variable | # Polymorphic Loci | Proportion Polymorphic | Average variable sites per variable locus |
| --- | --- | --- | --- | --- | --- | --- | --- | --- | --- |
| *Antrodiaetus* | 480 | 329,110 | 685.65 | 129-1248 | 8,389 | 2.55 | 476 | 0.99 | 17.62 |
| *Briggsus* | 292 | 172,562 | 590.97 | 144-1190 | 25,524 | 14.79 | 292 | 1.00 | 87.41 |
| *Centruroides* | 585 | 583,454 | 997.36 | 243-1164 | 7,660 | 1.31 | 527 | 0.90 | 14.54 |

Maximum-likelihood analyses demonstrated the utility of the arachnid-specific UCE probe set in resolving relationships among and within arachnid orders (Figs. 1 - 3). For the limited deep-level sample dataset (i.e., ‘2perArachnid’), bootstrap support was 100% for all nodes except the node uniting Scorpiones to Araneae + Pedipalpi (=Amblypygi + Thelyphonida) (Fig. 1). For the Araneae dataset, most nodes were fully supported (BS=100%), while two nodes within Entelegyne spiders had bootstrap values of 97% (Fig. 2). Within Opiliones, all nodes received bootstrap support ≥99% (Fig. 3), and within Scorpiones, all nodes were fully supported except for one node with bootstrap support of 69% (Fig. 3).

**Figure 1.**
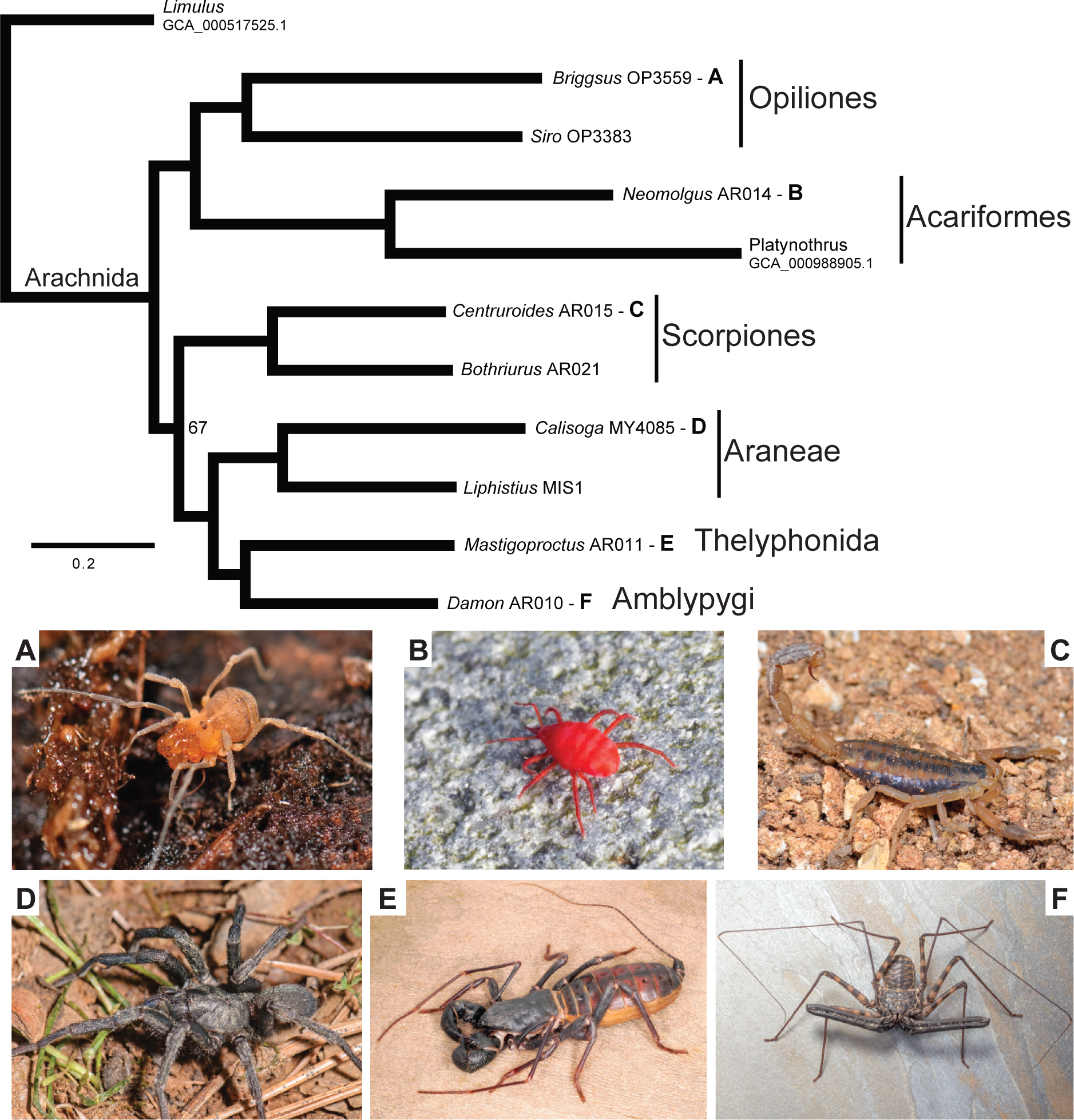
Maximum likelihood phylogeny based on 50% minimum taxon coverage “2perArachnid” dataset, including representative images of included arachnid orders. Nodes have 100% bootstrap support, unless otherwise indicated. Letters A-F correspond to representative images of included arachnid orders. Photo credit: M Hedin (A, C, D), M Erbland (B), WE Savary (E, F).

## Discussion

### Arachnid UCEs

The development of genetic markers for arachnids has lagged behind that of many groups due, in part, to the ancient divergences within the group (at least 400 MYA; Rehm *et al.* 2012; Rota-Stabelli *et al.* 2013) and the relatively few published arachnid genomic resources. Here, we demonstrate the utility of a UCE probe set targeting around 1000 loci that successfully works for all arachnid species tested (and likely also with close outgroups). Across all samples, which includes the deepest divergence within Arachnida (Regier *et al.* 2010; Sharma *et al.* 2014), we recovered 510 UCE loci with more than 49,362 parsimony-informative sites. For datasets within scorpions, spiders, and harvestmen, which each span root nodes estimated at hundreds of millions of years of divergence (Hedin *et al.* 2012; Bond *et al.* 2014; Sharma and Wheeler 2014), we recovered 749, 724, and 381 loci, respectively. We did not find a significant correlation between library pool size and average number of loci recovered (Fig. S1), and thus the lower number of loci recovered for Opiliones likely reflects their distant relationship to the taxonomic groups from which the probes were developed.

### Phylogenetic Utility

Recent phylogenomic datasets based on transcriptomes have provided better-resolved and well-supported phylogenies compared to prior morphological and genetic work using few loci (Hedin *et al.* 2012; Bond *et al.* 2014; Fernandez *et al.* 2014; Sharma *et al.* 2014; Sharma *et al.* 2015; Garrison *et al.* 2016). However, obtaining sequence coverage across taxa from transcriptomes requires high-quality RNA and a consistent expression pattern, which may be difficult to obtain for many non-model taxa. UCE sequence data have proven to be useful for resolving relationships at both deep and shallow levels (Crawford *et al.* 2012; Faircloth *et al.* 2012; McCormack *et al.* 2012; Faircloth *et al.* 2013; McCormack *et al.* 2013; Smith *et al.* 2014; Streicher *et al.* 2015; Meiklejohn *et al.* 2016), and can be obtained from museum specimens or other potentially degraded samples (McCormack *et al.* 2015).

Our reconstructed phylogenies based on UCEs demonstrate the promising utility of this probe set for collecting data to resolve relationships at a wide range of divergence levels, from the deepest bifurcations in the arachnid tree to shallower divergences within genera. In our limited deep level sample dataset (i.e., ‘2perArachnid’), we recovered well-supported monophyletic groups for all orders for which more than one individual was included (Fig. 1). Additionally, we obtained well-supported sister relationships between Thelyphonida and Amblypygi (Pedipalpi), and a Pedipalpi sister group to Araneae, consistent with relationships based on morphology, Sanger sequence data, and transcriptome data (Giribet *et al.* 2002; Shultz 2007; Regier *et al.* 2010; Sharma *et al.* 2014). The low support in our phylogeny for the placement of Scorpiones may be a result of our sparse sampling, or it may reflect the difficulty in determining the position of this group due to a possible rapid radiation deep in the arachnid tree of life (Sharma *et al.* 2014).

Phylogenetic analyses of the UCE data for our more densely sampled groups (Araneae, Opiliones, Scorpiones; Figs. 2 and 3) also recover topologies that are mostly congruent with recent transcriptome-based phylogenies (Hedin *et al.* 2012; Bond *et al.* 2014; Sharma *et al.* 2015; Garrison *et al.* 2016). Within Araneae (Fig. 2), many well-supported relationships and clades are recovered including Opisthothelae (Mygalomorphae + Araneomorphae), a split between the Atypoidea and Avicularioidea within mygalomorphs, and a split between Haplogynae and Entelegynae within araneomorphs. We also recover a well-supported Paleocribellate (*Hypochilus*) + Filistatidae (*Kukulcania*) sister group, a relationship only recently hypothesized based on transcriptome data (Bond *et al.* 2014). Relationships within Entelegynae using UCE data were not consistent with those recovered with transcriptome data (Bond *et al.* 2014; Garrison *et al.* 2016). However, there is disagreement in relationships between Araneoidea, RTA spiders, and related taxa in the different transcriptome datasets, and support values are relatively low at these nodes for both transcriptome and UCE data, indicating need for further sampling. Within Opiliones (Fig. 3), we recover monophyletic Palpatores (Eupnoi + Dyspnoi) and Laniatores, but we did not recover a monophyletic Phalangida (Palpatores + Laniatores) as recovered with transcriptomes (Hedin *et al.* 2012; Sharma *et al.* 2014) and a five gene dataset combined with morphological characters (Sharma and Giribet 2014). Within Scorpiones (Fig. 3), phylogenetic analyses of UCEs recover the same split between parvorders Buthida and Iurida as inferred from analyses of transcriptomes (Sharma *et al.* 2015). However, the well-supported position of *Bothriurus* deep within Iurida in our phylogeny, rather than as an early diverging species in Iurida based on transcriptome analyses (Sharma *et al.* 2015), highlights the need for further investigation using genome wide markers for this ancient group.

**Figure 2.**
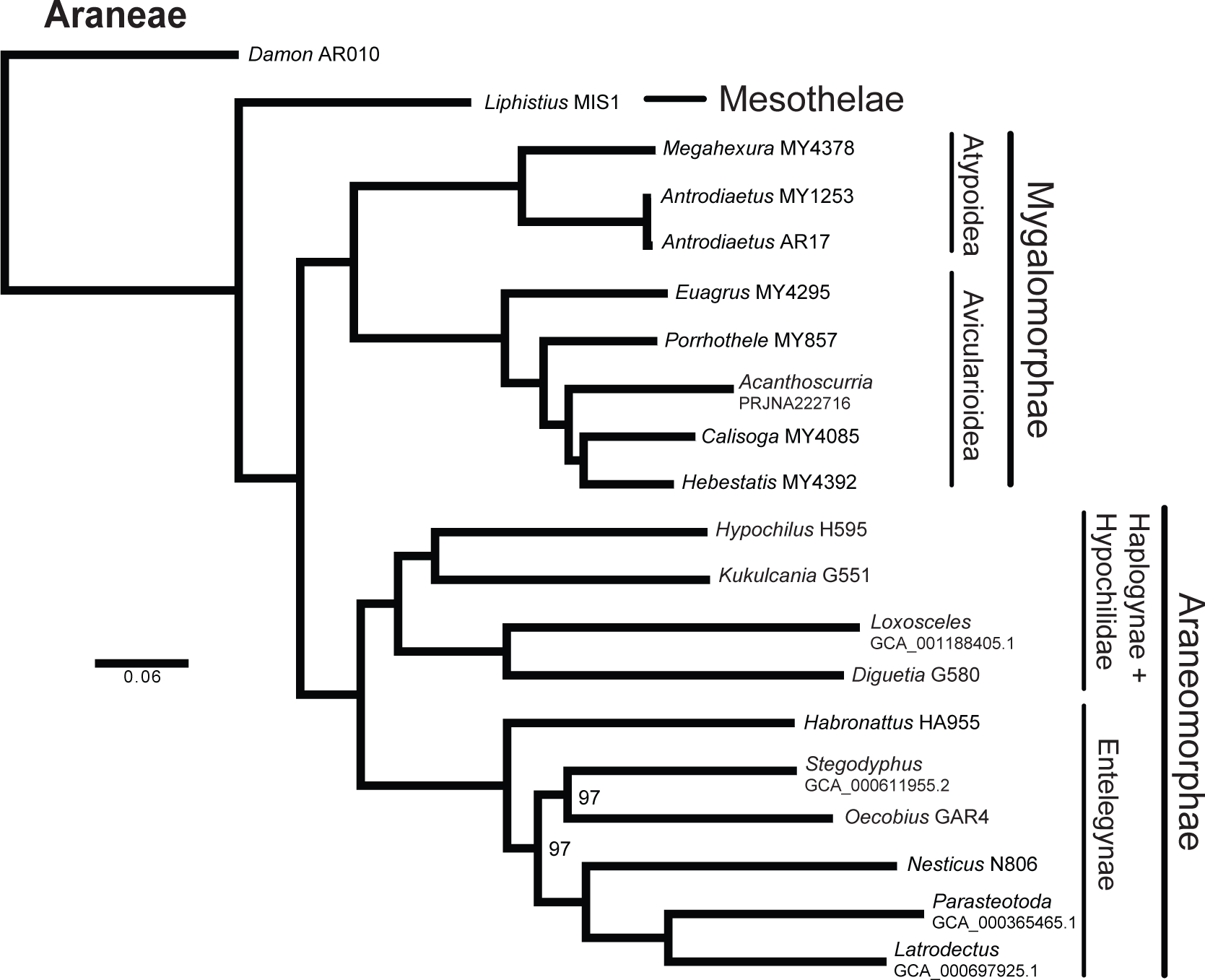
Maximum likelihood phylogeny based on 50% minimum taxon coverage dataset for 491 the Araneae+OUT dataset. Nodes have 100% bootstrap support, unless otherwise indicated.

Comparison of UCE sequences across close relatives within spiders, harvestmen, and scorpions indicate that our probe set will have applications beyond deep-level systematics and taxonomy. The UCE loci show promise for use in species-level phylogenetics and species delimitation because the flanking regions show increasing variability as distance from the core UCE increases (Fig. 3). UCEs enrichment produced 480 loci with more than 8,000 variable sites across two members of the *Antrodiaetus riversi* complex, 292 loci with more than 25,000 variable sites across two harvestmen species in the genus *Briggsus*, and 585 loci with more than 7,000 variable sites across an individual of the scorpion genus *Centruroides* and the published *Centruroides* genome (Table 3). These patterns of UCE variability are correlated with divergence in mitochondrial cytochrome oxidase I (Fig. 4). The proportion of polymorphic loci (minimum 0.9) and average variable sites per polymorphic locus (minimum 14) seen across these pairs is greater than those reported in the study of Smith *et al.* (2014; maximum 0.77 and 3.2, respectively), which demonstrated the utility of UCEs at shallow time scales for species-level phylogenetic and species delimitation analyses in birds. UCEs may be advantageous for shallow-level studies in Arachnida because other techniques, such as RADseq, are susceptible to locus dropout in arachnid species with relatively deep levels of divergence (Bryson *et al*. 2016; Derkarabetian *et al.* in press). Our UCE probe set will also provide a complementary/alternative resource to the recently developed spider anchored-enrichment loci, which were used to produce a comparable number of loci (455) and a well-resolved phylogeny for the North American tarantula genus *Aphonopelma* (Hamilton *et al*. 2016).

**Figure 3.**
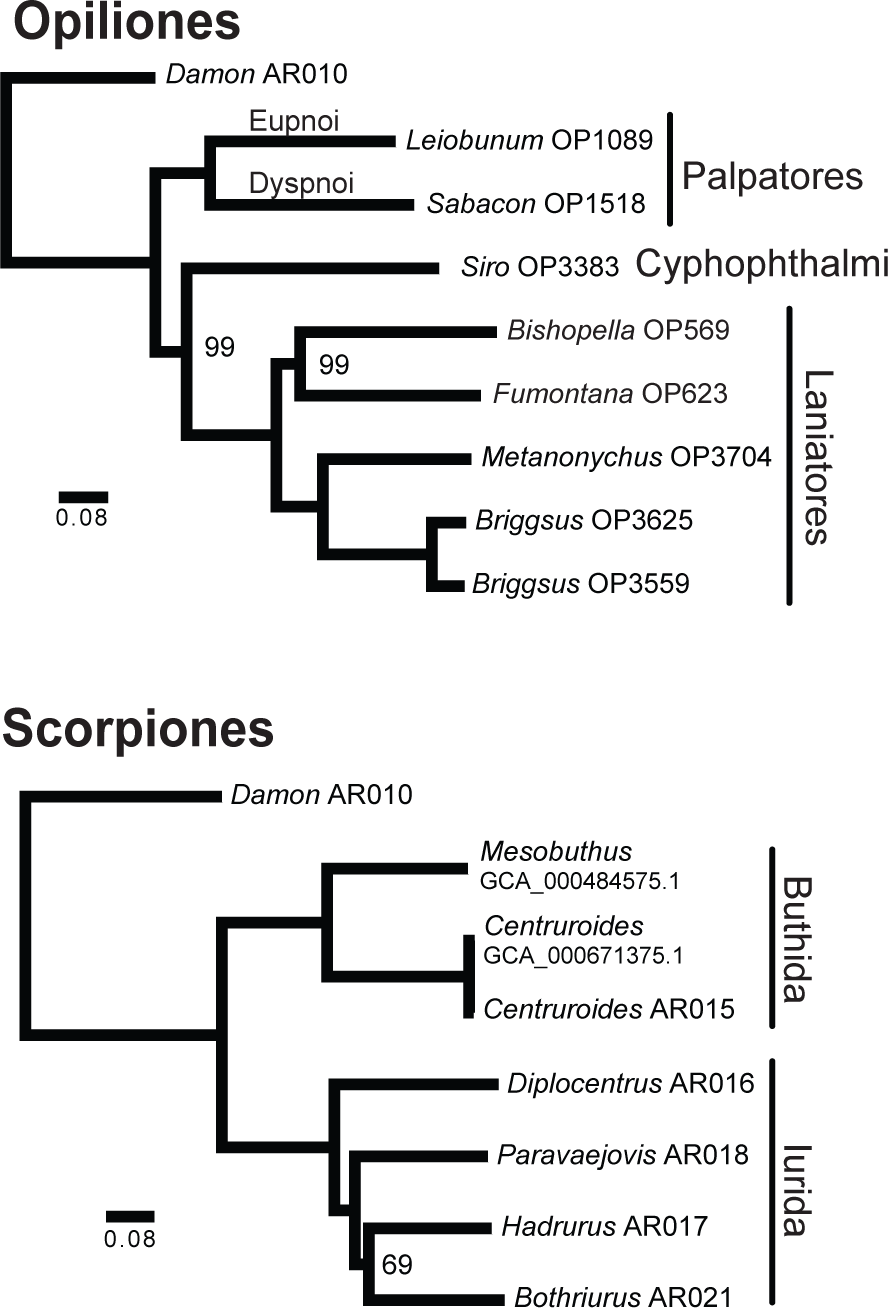
Maximum likelihood phylogenies based on 50% minimum taxon coverage dataset for the Opiliones+OUT and Scorpiones+OUT datasets. Nodes have 100% bootstrap support, unless otherwise indicated.

**Figure 4.**
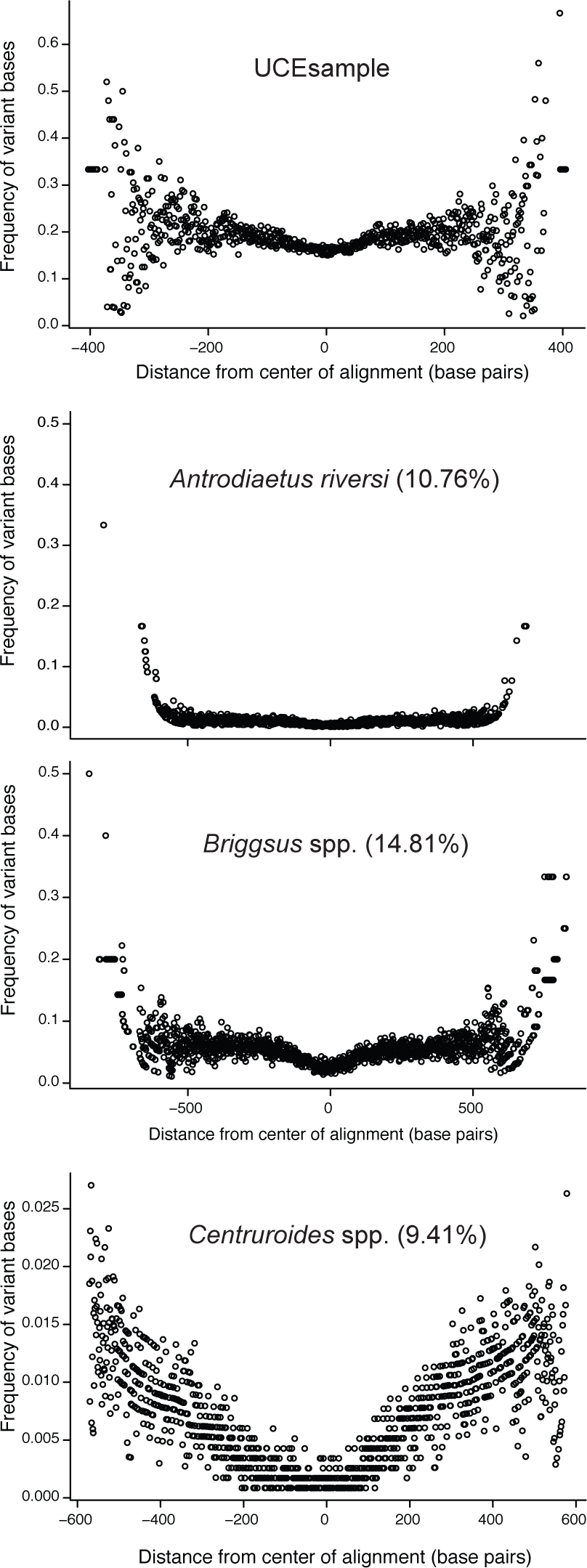
Variability increases as distance from core UCE increases. Data points with no variability were removed. For *Antrodiaetus* (turret spiders) and *Briggsus* (harvestmen) comparisons, the scale of x-axis is identical. Numbers in parentheses are uncorrected COI divergence values. In *Antrodiaetus*, an outlier was removed for better viewing. Cytochrome oxidase subunit I (COI) distances for *Centruroides* (bark scorpions) calculated from GenBank accessions AY995831.1 (*C. sculpturatus*) and AY995833.1 (*C. exilicauda*).

The arachnid-specific UCEs have utility and phylogenetic informativeness at all levels of Arachnida, spanning extremely ancient divergence times between orders (at least 400 MYA) to more recent congeneric divergences (<10 MYA for *Antrodiaetus*, Hedin *et al.* 2012). Thus, these markers will be an integral component of future comparative studies. Arachnids are an extremely diverse group, and yet it is estimated that less than half of arachnid species have been formally described (Chapman 2009). Species delimitation increasingly relies on phylogenomic data that can be sampled consistently across diverse taxa (Leaché *et al*. 2014; Smith *et al*., 2014; Rannala 2015; Harvey *et al*. 2016). The presence of UCEs across Arachnida makes these a valuable resource for discovery of new species and for inferring phylogenetic relationships in understudied arachnid groups.

## Acknowledgements

For help during lab work we thank Carl Oliveros and Rich Harrington. John Klicka provided other assistance. Tim Burkhart, Ken MacNeil, Peter Clausen, and Cor Vink provided assistance with specimen collection. Live specimen photos were provided by Mardon Erbland and Warren E. Savary. Funding for this research was provided by NSF grant DEB-1354558 to MH. Startup funds from LSU and DEB-1242260 to BCF supported computational portions of this work.

## Data Availability

Raw sequence reads are available in the NCBI Short Read Archive accession SRP078995 (BioProject ID #PRJNA328972), and untrimmed contigs have been given accessions ######-#####. Trimmed alignments used for phylogenetic analyses are available in Dryad (doi: ######).

## Author Contributions

All authors contributed to the conception and design of the experiments; JS, SD, MH, RWB contributed samples; JS, SD, RWB, BCF conducted lab work; JS, SD, BCF analyzed data; all authors edited and commented on the manuscript.

## References

Alexander RP, Fang G, Rozowsky J, Snyder M, Gerstein MB (2010) Annotating non-coding regions of the genome. Nature Reviews Genetics, 11, 559–571.

Bejerano G, Pheasant M, Makunin I, Stephen S, Kent WJ, Mattick JS, Haussler D (2004) Ultraconserved elements in the human genome. Science, 304, 1321–1325.

Bond JE, Garrison NL, Hamilton CA, Godwin RL, Hedin M, Agnarsson I (2014) Phylogenomics resolves a spider backbone phylogeny and rejects a prevailing paradigm for orb web evolution. Current Biology, 24, 1765–1771. doi.org/10.1016/j.cub.2014.06.034.

Bryson RW, Savary WE, Zellmer AJ, Bury RB, McCormack JE (2016) Genomic data reveal ancient microendemism in forest scorpions across the California Floristic Province. Molecular Ecology, doi: 10.1111/mec.13707.

Castresana J (2000) Selection of conserved blocks from multiple alignments for their use in phylogenetic analysis. Molecular Biology and Evolution, 17, 540–552.

Chapman AD (2009) Numbers of Living Species in Australia and the World, (2nd edn), Australian Biodiversity Information Services.

Crawford NG, Faircloth BC, McCormack JE, Brumfield RT, Winker K, Glenn TC (2012) More than 1000 ultraconserved elements provide evidence that turtles are the sister group of archosaurs. Biology Letters, 8, 783–786. doi:10.1098/rsbl.2012.0331.

Derkarabetian S, Burns M, Starrett J, Hedin M. (2016) Population genomic evidence for multiple refugia in montane-restricted harvestmen (Arachnida, Opiliones, Sclerobunus robustus) from the southwestern United States. Molecular Ecology, In Press.

Derti A, Roth FP, Church GM, Wu C (2006) Mammalian ultraconserved elements are strongly depleted among segmental duplications and copy number variants. Nature Genetics, 38, 1216–1220. doi:10.1038/ng1888

Dunlop JA (1996) A trigonotarbid arachnid from the Upper Silurian of Shropshire. Palaeontology, 39, 605–614.

Dunlop JA, Tetlie OE, Prendini L (2008) Reinterpretation of the Silurian scorpion *Proscorpius osborni* (Whitfield): integrating data from Palaeozoic and Recent scorpions. Palaeontology, 51, 303–320.

Faircloth BC (2016) Identifying conserved genomic elements and designing universal probe sets to enrich them. Molecular Ecology Resources, submitted.

Faircloth BC, Branstetter MG, White ND, Brady SG (2015) Target enrichment of ultraconserved elements from arthropods provides a genomic perspective on relationships among Hymenoptera. Molecular Ecology Resources, 15, 489–501. doi: 10.1111/1755–0998.12328.

Faircloth BC, McCormack JE, Crawford NG, Harvey MG, Brumfield RT, Glenn TC (2012) Ultraconserved elements anchor thousands of genetic markers spanning multiple evolutionary timescales. Systematic Biology, 61, 717–726. DOI:10.1093/sysbio/sys004.

Faircloth BC, Sorenson L, Santini F, Alfaro ME (2013) A phylogenomic perspective on the radiation of ray-finned fishes based upon targeted sequencing of ultraconserved elements (UCEs). PLoS ONE 8, e65923. doi:10.1371/journal.pone.0065923.

Fernández R, Hormiga G, Giribet G (2014) Phylogenomic analysis of spiders reveals nonmonophyly of orb weavers. Current Biology, 24, 1772–1777. doi.org/10.1016/j.cub.2014.06.035.

Garrsion NL, Rodriguez J, Agnarsson I, Coddington JA, Griswold CE, Hamilton CA, Hedin M, Kocot KM, Ledford JM, Bond JE (2016) Spider phylogenomics: untangling the spider tree of life. PeerJ, 4:e1719; DOI 10.7717/peerj.1719.

Gilbert PS, Chang J, Pan C, Sobel EM, Sinsheimer JS, Faircloth BC, Alfaro ME (2015) Genome-wide ultraconserved elements exhibit higher phylogenetic informativeness than traditional gene markers in percomorph fishes. Molecular Phylogenetics and Evolution, 92, 140–146. doi.org/10.1016/j.ympev.2015.05.027.

Giribet G, Edgecombe GD, Wheeler WC, Babbitt C (2002) Phylogeny and systematic position of opiliones: A combined analysis of chelicerate relationships using morphological and molecular data1. Cladistics, 18, 5–70.

Glenn TC, Nilsen R, Kieran TJ, Finger JW, Pierson TW, Bentley KE, et al. (2016) Adapterama I: Universal stubs and primers for thousands of dual-indexed Illumina libraries (iTru & iNext). bioRxiv, doi: 10.1101/049114.

Grabherr MG, Haas BJ, Yassour M, Levin JZ, Thompson DA, Amit I, Adiconis X, Fan L, Raychowdhury R, Zeng Q, Chen Z, Mauceli E, Hacohen N, Gnirke A, Rhind N, di Palma F, Birren BW, Nusbaum C, Lindblad-Toh K, Friedman N, Regev A (2011) Full-length transcriptome assembly from RNA-Seq data without a reference genome. Nature Biotechnology, 29, 644–652.

Hamilton CA, Hendrixson BE, Bond JE (2016) Taxonomic revision of the tarantula genus *Aphonopelma* Pocock, 1901 (Araneae, Mygalomorphae, Theraphosidae) within the United States. ZooKeys, 560, 1.

Harvey MG, Smith BT, Glenn TC, Faircloth BC, Brumfield RT (2016) Sequence Capture Versus Restriction Site Associated DNA Sequencing for Shallow Systematics. Systematic Biology, doi: 10.1093/sysbio/syw036.

Harvey MS (2002) The neglected cousins: What do we know about the smaller arachnid orders? Journal of Arachnology, 30, 357–372.

Hedin M, Starrett J, Akhter S, Schönhofer AL, Shultz JW (2012) Phylogenomic resolution of paleozoic divergences in harvestmen (Arachnida, Opiliones) via analysis of next-generation transcriptome data. PLoS ONE, 7, e42888. doi:10.1371/journal.pone.0042888.

Hedin M, Starrett J, Hayashi C (2013) Crossing the uncrossable: novel trans-valley biogeographic patterns revealed in the genetic history of low-dispersal mygalomorph spiders (Antrodiaetidae, Antrodiaetus) from California. Molecular Ecology, 22, 508–526.

Jeram AJ, Selden PA, Edwards D (1990) Land animals in the Silurian: arachnids and myriapods from the Shropshire, England. Science, 250, 658–661.

Katoh K, Standley DM (2013) MAFFT multiple sequence alignment software version 7: improvements in performance and usability. Molecular Biology and Evolution, 30, 772–780.

Leaché AD, Fujita MK, Minin VN, Bouckaert RR (2014) Species Delimitation using genome-wide SNP data. Systematic Biology. 63, 534–542. doi:10.1093/sysbio/syu018.

Masta SE, Longhorn SJ, Boore JL (2009) Arachnid relationships based on mitochondrial genomes: asymmetric nucleotide and amino acid bias affects phylogenetic analyses. Molecular Phylogenetics and Evolution, 50, 117–128.

Laurie M (1899) On a Silurian scorpion and some additional eurypterid remains from the Pentland Hills. Transactions of the Royal Society of Edinburgh, 39, 575–590.

Manthey JD, Campillo LC, Burns KJ, Moyle RG (2016) Comparison of target-capture and restriction-site associated DNA sequencing for phylogenomics: a test in cardinalid tanagers (Aves, Genus: Piranga). Systematic biology, 10.1093/sysbio/syw005.

McCormack JE, Faircloth BC, Crawford NG, Gowaty PA, Brumfield RT, Glenn TC (2012) Ultraconserved elements are novel phylogenomic markers that resolve placental mammal phylogeny when combined with species-tree analysis. Genome Research, 22, 746–754.

McCormack JE, Harvey MG, Faircloth BC, Crawford NG, Glenn TC, Brumfield RT (2013) A phylogeny of birds based on over 1,500 loci collected by target enrichment and high-throughput sequencing. PLoS One, 8, e54848.

McCormack JE, Tsai WLE, Faircloth (2015) Sequence capture of ultraconserved elements from bird museum specimens. Molecular Ecology Resources, 1–15. doi: 10.1111/1755–0998.12466.

Meiklejohn KA, Faircloth BC, Glenn TC, Kimball RT, Braun EL (2016) Analysis of a rapid evolutionary radiation using ultraconserved elements (UCEs): Evidence for a bias in some multispecies coalescent methods. Systematic Biology, 65, 612–627.

Rannala B (2015) The art and science of species delimitation. Current Zoology. 61, 846–853.

Regier JC, Shultz JW, Zwick A, Hussey A, Ball B, Wetzer R, Martin JW, Cunningham CW (2010) Arthropod relationships revealed by phylogenomic analysis of nuclear protein-coding sequences. Nature, 463, 1079–1083. doi:10.1038/nature08742.

Rehm P, Pick C, Borner J, Markl J, Burmester T (2012) The diversity and evolution of chelicerate hemocyanins. BMC Evolutionary Biology, 12, 19.

Rota-Stabelli O, Daley AC, Pisani D (2013) Molecular timetrees reveal a Cambrian colonization of land and a new scenario for ecdysozoan evolution. Current Biology, 23, 392–398.

Sharma PP, Giribet G (2014) A revised dated phylogeny of the arachnid order Opiliones. Frontiers in Genetics, 5, 1–13. doi:10.3389/fgene.2014.00255.

Sharma PP, Fernández R, Esposito LA, González-Santillán E, Monod L (2015) Phylogenomic resolution of scorpions reveals multilevel discordance with morphological phylogenetic signal. Proceedings of the Royal Society of London B, 282, 20142953. doi.org/10.1098/rspb.2014.2953.

Sharma PP, Kaluziak ST, Pérez-Porro AR, González VL, Hormiga G, Wheeler WC, Giribet G (2014) Phylogenomic interrogation of arachnida reveals systematic conflicts in phylogenetic signal. Molecular Biology and Evolution, 31, 2963–2984. doi:10.1093/molbev/msu235.

Shultz JW (2007) A phylogenetic analysis of the arachnid orders based on morphological characters. Zoological Journal of the Linnean Society, 150, 221–265. DOI:10.1093/sysbio/syt061.

Smith BT, Harvey MG, Faircloth BC, Glenn TC, Brumfield RT (2014) Target capture and massively parallel sequencing of ultraconserved elements for comparative studies at shallow evolutionary time scales. Systematic Biology, 63, 83–95.

Stamatakis A, Hoover P, Rougemont J (2008) A rapid bootstrap algorithm for the RAxML web servers. Systematic biology, 57, 758–771.

Stamatakis A (2014) RAxML version 8: a tool for phylogenetic analysis and post-analysis of large phylogenies. Bioinformatics, 30, 1312–1313.

Streicher JW, Schulte II JA, Weins JJ (2016) How should genes and taxa be sampled for phylogenomic analyses with missing data? An empirical study in Iguanian lizards. Systematic Biology, 65, 128–145. DOI:10.1093/sysbio/syv058.

Talavera G, Castresana J (2007) Improvement of phylogenies after removing divergent and ambiguously aligned blocks from protein sequence alignments. Systematic biology, 56, 564–577.

Wheeler WC, Hayashi CY (1998) The phylogeny of the extant chelicerate orders. Cladistics, 14, 173–192.

Zhang Z-Q (2011) Phylum Arthropoda von Siebold, 1848 In: Zhang, Z.-Q. (Ed.) Animal biodiversity: An outline of higher-level classification and survey of taxonomic richness. Zootaxa, 3148, 99–103.

